# First Draft Genome of a Brazilian Atlantic Rainforest Burseraceae reveals commercially-promising genes involved in terpenic oleoresins synthesis

**DOI:** 10.1101/467720

**Authors:** Luana Ferreira Afonso, Danielle Amaral, Marcela Uliano-Silva, André Luiz Quintanilha Torres, Daniel Reis Simas, Mauro de Freitas Rebelo

**Affiliations:** Bio Bureau Biotechnology, Rio de Janeiro, RJ, Brazil; Biophysics Institute Carlos Chagas Filho, Federal University of Rio de Janeiro, RJ, Brazil; Leibniz Institute for Zoo and Wildlife Research (IZW), Berlin, Germany; Berlin Center for Genomics in Biodiversity Research (BeGenDiv), Berlin, Germany

**Author notes:** authors contributed equally (Luana Ferreira Afonso, Danielle Amaral).

## Abstract

**Background:** *Protium* species produce abundant aromatic oleoresins composed mainly of different types of terpenes, which are highly sought after by the flavor and fragrance industry.

**Results:** Here we present (i) the first draft genome of an endemic tree of the Brazil’s Atlantic Rainforest (Mata Atlântica), *Protium kleinii* Cuatrec., (ii) a first characterization of its genes involved in the terpene pathways, and (iii) the composition of the resin’s volatile fraction. The de novo draft genome was assembled using Illumina paired-end-only data, totalizing 407 Mb in size present in 229,912 scaffolds. The N50 is 2.60 Kb and the longest scaffold is 52.26 Kb. Despite its fragmentation, we were able to infer 53,538 gene models of which 5,434 were complete. The draft genome of P. kleinii presents 76.67 % (62.01 % complete and 14.66 % partial) of plant-core BUSCO genes. InterProScan was able to assign at least one Gene Ontology annotation and one Pfam domain for 13,629 and 26,469 sequences, respectively. We were able to identify 116 enzymes involved in terpene biosynthesis, such as monoterpenes *α*-terpineol, 1,8-cineole, geraniol, (+)-neomenthol and (+)-(R)-limonene. Through the phylogenetic analysis of the Terpene Synthases gene family, three candidates of limonene synthase were identified. Chemical analysis of the resin’s volatile fraction identified four monoterpenes: terpinolene, limonene, *α*-pinene and *α*-phellandrene.

**Conclusion:** These results provide resources for further studies to identify the molecular bases of the main aroma compounds and new biotechnological approaches to their production.

## 1. INTRODUCTION

Among the secondary metabolites that plants produce, terpenes are the largest class, comprising more than 40,000 structurally different substances [1]. They play important roles as pollinator attractants, antimicrobials, antiherbivores, allelopathics, hormones, vitamins and membrane stabilizers [2]. Terpenes are also notable for their biotechnological applications in the food, cosmetic and pharmaceutical industries as flavors, fragrances, colorants, and medications, with demand growing in recent years [3, 4]. Protium species are known to produce more than one hundred terpenes [5]. The volatile fraction of their oleoresins is composed mainly of mono- and sesqui-terpenes while the solid fraction is composed of triterpenes [6]. Terpene production is catalyzed by enzymes encoded by the extremely diverse Terpene Synthase (TPS) family of genes [7, 8]. The monoterpene variety seen in Protium is thought to reflect this gene diversity [9], oleoresin modification over time [5] and the multiple catalytic functions observed in monoterpene biosynthesis [10]. Genome characterization is one of the most informative descriptions of a species’ biology [11]. It can contribute to a better understanding of metabolic pathways involved in the production of several metabolite classes and also identify the molecular basis, potentially illuminating new biotechnological approaches to their production [12]. Here we present a comprehensive analysis of the draft genome of the *Protium kleinii* Cuatrec. (Burseraceae Family).

## 2. EXAMPLES OF ARTICLE COMPONENTS

The sections below show examples of different article components.

## 3. RESULTS

### A. Genome assembly and gene content

This study aimed to achieve a de novo assembly of the first genome of a species belonging to the *Protium* genus with sufficient precision for gene identification. The Illumina run generated 525,305,928 raw reads and 99.08% of reads passed the cleaning steps and were used for assembly with SOAPdenovo. K-mer analysis (31 bp in size) estimated the genome size as 445 Mb with 1.15% heterozygosity (Fig. 1). The draft genome was assembled in 229,912 scaffolds with a N50 of 2.6 Kb and a final size of 407 Mb [Table 1]. The GC content was estimated to be 33.22% with 41.70% repetitive elements [Additional file 1: Table S1]. The maximum scaffold length generated was 52.26 Kb. It is not necessary to place figures and tables at the back of the manuscript. Figures and tables should be sized as they are to appear in the final article. Do not include a separate list of figure captions and table titles.

**Fig. 1.**
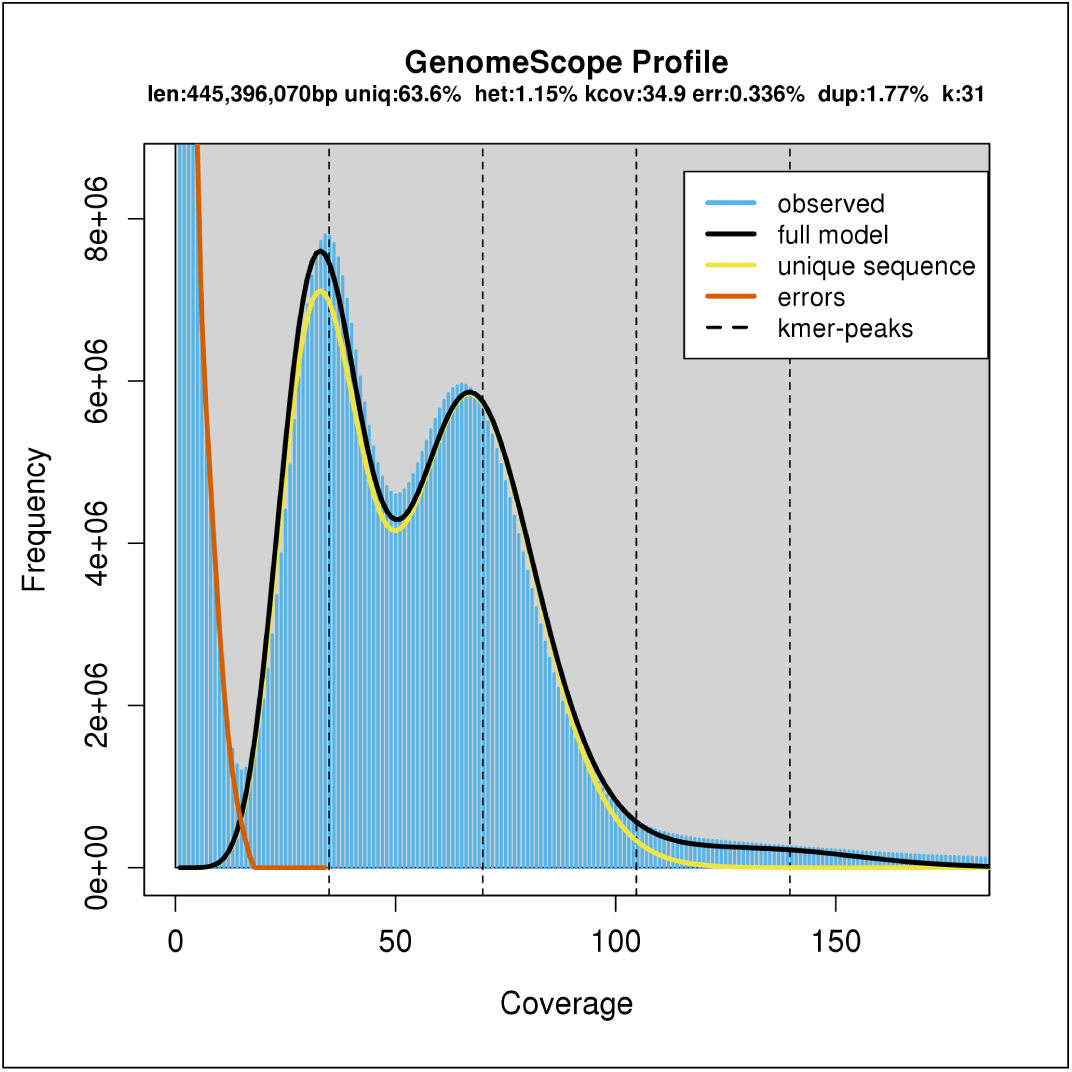
K-mer distribution of P. kleinii Illumina DNA reads.

**Table 1.** Assembly statistics for the draft genome of *P. kleinii*

| Parameter | Value |
| --- | --- |
| Estimated genome size by k-mer analysis, bp | 445,396,070 |
| Total size of assembled genome, Mb | 407.28 |
| Number of scaffolds | 229,912 |
| Number of contigs | 563,793 |
| Scaffold N50, Kb | 2.60 |
| Maximum scaffold length, Kb | 52.26 |
| Percentage of genome in scaffolds > 50 Kb | 0.03 % |
| Masked percentage of total genome | 41.76 % |

Scaffolds were evaluated for gene content with gVolante, which found 76.67% (partial or complete) of the 1,440 plant core genes [Additional file 2: Table S2]. In this first assembly, we have predicted 57,538 gene models, which were then annotated using different databases. BLASTp identified 29,995 (52.10%) hits against the UniProtKB/Swiss-Prot database and 12,952 (22.50%) using NCBI non-redundant databases [Additional file 3: Table S3]. Using the BLASTp results against the UniProt and perl scripts [13] we showed that this first assembly generated 5,434 gene models that are represented by nearly full-length transcripts, with >80% of alignment coverage once BLASTed against a Swiss-Prot database sequence. InterProScan analysis with all protein sequences was able to assign at least one Gene Ontology (GO) term annotation and one Pfam domain to 13,629 (23.70%) and 26,469 (46.00%) sequences, respectively, while KAAS (KEGG Automatic Annotation Server Version 2.1; [14]) assigned 7,477 (13.00%) KEGG Orthology (KO) to the putative proteins of *P. kleinii* [Table 2; Additional file 4: Table S4].

**Table 2.** Summary of gene prediction and general functional annotation of *P. kleinii*

| Parameter | Value |
| --- | --- |
| Total number of genes | 57,538 |
| Total number of exons | 193,945 |
| Total number of proteins | 57,538 |
| Average protein size | 221 aa |
| Number of proteins BLAST hits* with Uniprot | 29,995 |
| Number of proteins BLAST hits* with NR NCBI<sup>a</sup> | 12,952 |
| Number of proteins InterProScan hits with Pfam | 26,469 |
| Number of proteins with KO assigned by KEGG | 7,477 |

\* all considered hits had a maximum E-value of 1e-05
<sup>a</sup> no hits with Uniprot

The complete InterProScan analyses with the default threshold revealed significant similarities with at least one protein signatures database in 44,616 (77.50 %) of the putative protein sequences of *P. kleinii*. It was possible to retrieve 35,310 GO terms comprising the three categories of biological process, molecular function, and cellular component [Additional file 5: Table S5]. Molecular function retrieved the majority of GO annotation assignments (22,551), followed by biological process (10,195) and cellular component (2,602) [Additional file 6: Figure S1].

### B. Comparative genomic analysis

*P. kleinii* gene models were analyzed together with the gene sets of *Citrus sinensis* (NCBI ID: 10702; [15], *Arabdopsis thaliana* (Ensembl ID: TAIR10; [16], *Glycine max* (Ensembl ID: Glycine max v2.0; [17], *Vitis vinifera* (Ensembl ID: IGGP12x; [18], and the monocotyledone *Oryza sativa* (ASM465v1; [19], in order to find shared genes and specific expansions and retractions of *P. kleinii* [Fig. 2]. This analysis yielded 17,204 (66.50%) groups of orthologue-containing sequences of two or more species. Within these clusters, 8,456 have orthologues of all six species, where 581 comprise single-copy genes [Additional file 7: Table S6]. Most were assigned to GO terms related to essential processes such as DNA replication, transcription and translation machinery, cell cycle control, cell signaling, metabolism and transport. A summary containing the number of clusters by GO categories is available in Additional file 8: Table S7. Although we have used the protein dataset of closely related-species of plants such as *C. sinensis* and the model organism, *A. thaliana* [20], 1,795 clusters were unique to *P. kleinii*. Of these, 1,214 (67.60%) had no protein with a significant hit against the UniProt database and, hence, were classified as “unknown function” [Additional file 9: Table S8]. Among those that could be annotated, 36 clusters comprise 1,333 sequences with elements commonly found in viral polyproteins. This comparative analysis, also allowed us to identify putative cases of gene loss in *P. kleinii*. Of all clusters, 1,300 had at least one orthologue of each species, except for *P. kleinii* [Additional file 9: Table S8]. This analysis also revealed 26,497 singletons [Additional file 10: Table S9].

**Fig. 2.**
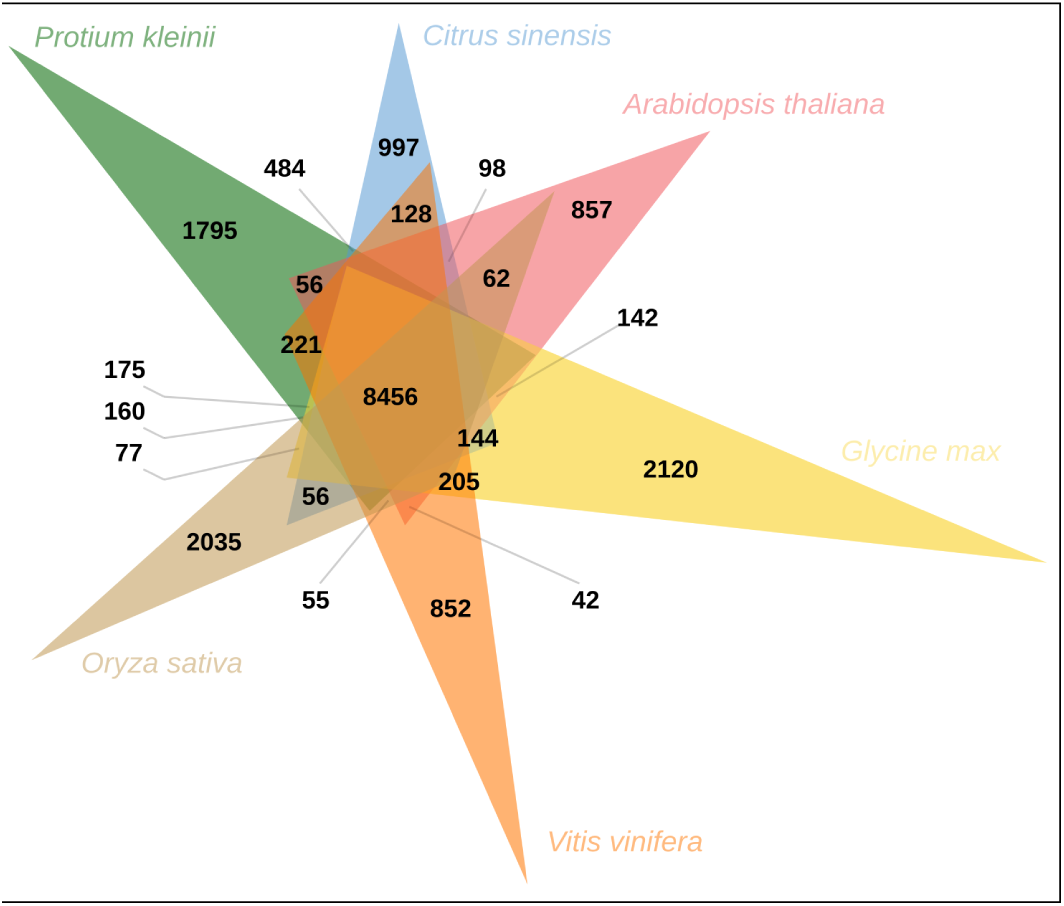
OrthoVenn diagram presenting the number of clustered genes in *P. kleinii* and five other plant species. *P. kleinii* area is colored in green, *C. sinensis* - blue, *A thaliana* - pink, *G. max* - yellow, *V. vinifera* - orange and *O. sativa* - light brow. Shared orthologous among all species are contained in 8,456 clusters. Putative specific gene expansions of *P. kleinii* are comprised in 1,795 clusters.

### C. *P. kleinii* essential oil analysis

In order to investigate the chemical profile of the essential oil extracted from the resin of *P. kleinii* (the yield was 5.00%), Gas-phase Chromatography coupled to Mass Spectrometry (GC-MS) was performed. This chemical analysis revealed the presence of terpinolene (79.66%); limonene (8.72%); *α*-pinene (5.55%) and *α*-phellandrene (1.57%) [Additional file 11: Table S10] Together these four compounds comprised 95.50% of the extracted oil. The chromatogram obtained by GC-MS analysis and mass spectrometry spectra are available in additional file 12: Documentation 1. We believe this is the first published report detailing the composition of the volatile fraction of *P. kleinii* resin.

### D. *In silico* identification of Terpene Biosynthesis Pathway genes

In a first attempt to identify genes of *P. kleinii* involved in the molecular pathways of terpene biosynthesis, we used the Kyoto Encyclopedia of Genes and Genomes database (KEGG; [21]) to search for enzymes assigned to these processes. We were able to map 115 enzymes to the four terpene pathways. Seventy belonged to the terpenoid backbone pathway (map00900) [Additional file 13: Figure S2], 13 to monoterpene (map00902) [Additional file 14: Figure S3], 22 to diterpene (map00904) [Additional file 15: Figure S4] and 10 to tri- and sesquiterpenes (map00909) [Additional file 16: Figure S5]. And limonene synthase was found employing a phylogenetic methodology. The complete set of enzymes of the plastidial methylerythritol phosphate (MEP) pathway and five of six enzymes primarily from the cytosolic mevalonic acid (MVA) pathway found in the other four rosids were mapped to *P. kleinii* using the KEGG database. From this terpenoid backbone biosynthesis pathway, the other three terpenoids biosynthesis pathways (maps 00902, 00904, 00909) are derived [Additional file 13: Figure S2]. From the 31 KO groups related to monoterpene biosynthesis, we were able to map three TPS into *α*-terpineol and 1,8-cineole producing enzymes (group K07385) and two into geranylpyrophosphate (GPP) to geraniol transformation enzymes (group K20979). We also found eight (+)-neomenthol dehydrogenases (group K15095), that are depicted in [Additional file 14: Figure S3]. As a second strategy to confirm the integrity of the monoterpene pathway, we searched for missing orthologues in the scaffold of *P. kleinii*. Although we had been using the protein sequence of the closest plant orthologue available in the KEGG database [Additional file 17: Table S11] in the query, Exonerate did not succeed in finding any of the absent genes [Additional file 14: Figure S3]. Due to their importance in the monoterpene production [5, 8], we searched all TPS gene families in the *P. kleinii* genome taking both conserved domains as references (PF01397.20 and PF03936.15; [22]. With this approach, we managed to recover 68 TPS of *P. kleinii*, where 20 were considered full length and 48 were partially predicted [Additional File 18: Table S12]. This enabled us to assess the evolution of the whole TPS genes family in *P. kleinii*. Of the nine query TPS, only limonene synthase of C. sinensis (KEGG accession number: 102607029) nested with three TPS of *P. kleinii* (scaffold100846.g17599.t1; scaffold187198.g29710.t1 and scaffold3882.g770.t1) and one TPS of P. decandrum (NCBI accession number: KC881151.1) in a clade with a reliable value of bootstrap (94% of bootstrap) (purple bar; Figure 3). In spite of the low values of bootstrap, eight TPS of *P. kleinii* TPS were grouped with TPS (fragments 2) belonging to the clades identified as C2 (66% of bootstrap), C3 (100% of bootstrap) and C4/C5 (100% of bootstrap) by authors Zapata and Fine [9] (yellow bar; Figure 3). The assembly of the *P. kleinii* genome allowed us to identify at least 60 putative TPS that were never previously studied (green bar; [Fig. 3]).

**Fig. 3.**
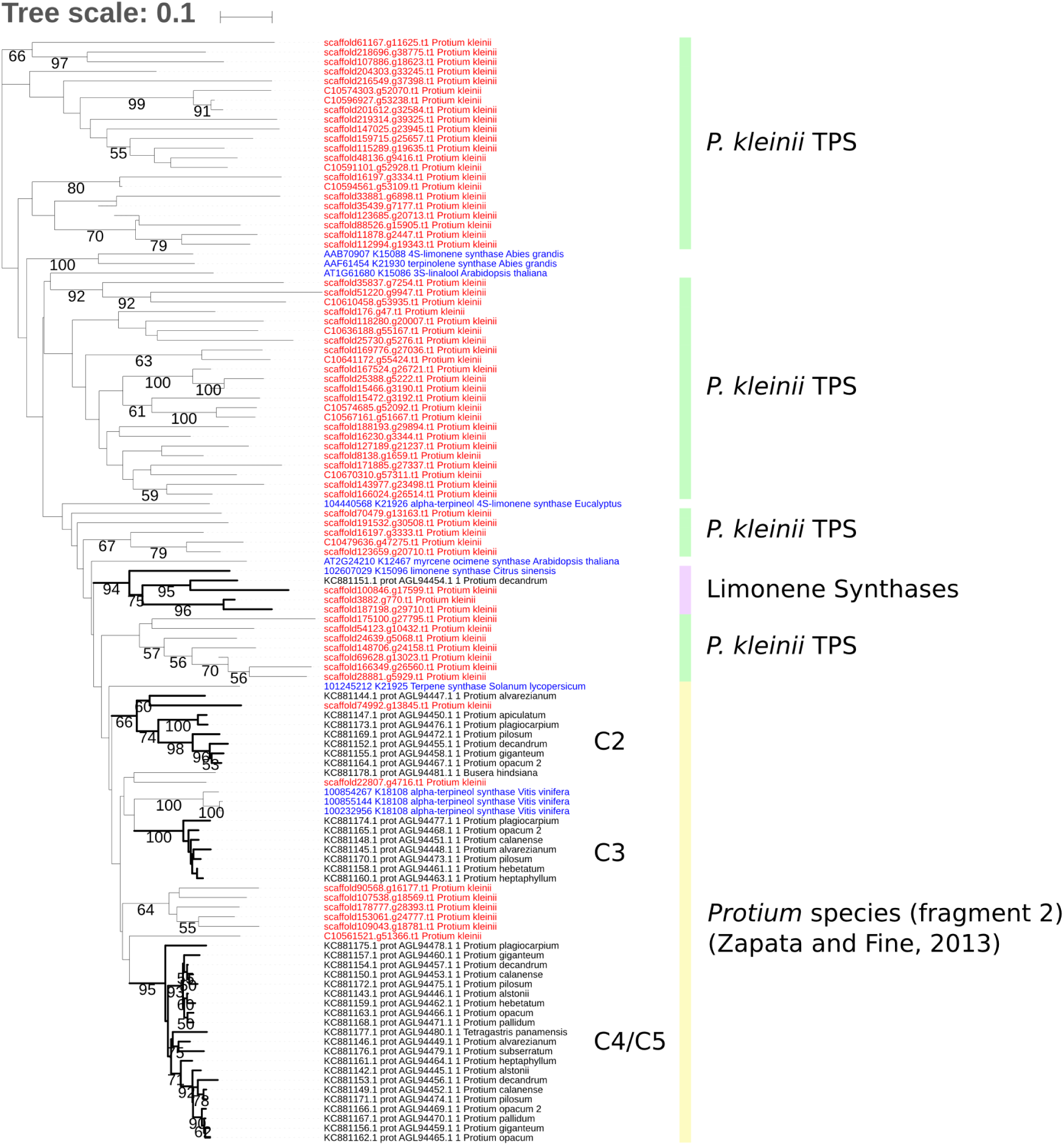
Phylogenetic tree of the TPS gene family.Green bars shows the clades containing paralogues of *P. kleinii*. Limonene synthase orthologues are highlighted with a purple bar. Yellow bar shows the clades C2, C3 and C4/C5 described in Zapata and Fine [9]. TPS recovered from the genome of *P. kleini*i are colored in red, queries accessed in the KEGG database are in blue and sequences retrieved (fragment 2) from Zapata and Fine [9] are in black. Only bootstrap higher than 50% are shown.

## 4. DISCUSSION

In this work we present the first genome characterization of *P. kleinii*, a Burseraceae from the Brazilian Atlantic Rainforest (Mata Atlântica). After consulting the Genomes Online Database (GOLD) [23] we believe we are the first to contribute the genome of a tree endemic to Brazil that occurs in the Mata Atlântica. Compounds of its resin essential oil demonstrate antinociceptive and anti-inflammatory effects [24, 25]. The plant biodiversity of Brazilian biomes is considered the most diverse in the world [26]. Sequencing the genomes of these species for conservation purposes and for biotechnology applications represents an enormous challenge. Just to assemble the complete genome of a single plant poses daunting challenges from both technological and cost perspectives. Draft genomes can offer a cost-effective alternative for the prediction of genes that could be employed in the drug discovery process, by the cosmetics industry, and in flora conservation efforts [5, 25] This study demonstrates how the assembled genome of *P.kleinii* could allow us to begin to investigate a genus that has species able to produce abundant aromatic oleoresins featuring different types of terpenes [5, 25], identifying specific genes and corresponding molecular pathways worthy of further study. The assembled genome has a total size of 407 Mb, which is comparable to the predicted genome size (445 Mb) [Figure 1] and is in accordance with the genome size of other rosids such as *C. medica* (404,9 Mb, [27] and *V. vinifera* (475 Mb, [18]. The low N50 in this work can be attributed to two main factors (i) the use of only paired-end Illumina data for genome assembly, and (ii) the high level of heterozygosity (1.15%) encountered with *P. kleinii*. The N50 is an important metric used to describe the contiguity and completeness of an assembled genome. Generally, the greater the N50, the better the genome assembly and the more accurate the prediction of complete genes and gene copies [28]. Nevertheless, even with this fragmented first draft, we were able to predict 76.67% of the genes – 62.01% complete genes and 14.66% partial genes – considered to be commonly shared among plants. Our AUGUSTUS gene prediction resulted in 57,538 gene models, which is higher than the gene dataset of other Sapindales such as *C. medica* (32,579 genes, [27], *C. unshiu* (29,024, [29], *C. sinensis* (29,445, [15] and even for other rosids such as *V. vinifera* (30,434, [18] and *A. thaliana* (25,498, The *Arabidopsis* Genome [30]. The higher number of genes in *P. kleini*i can be attributed to the prediction of cleaved genes and splitting loci [31, 32] because we did not use any long DNA reads to build long scaffolds and transcriptome data to support the gene prediction analysis. Despite fragmentation and overestimation of gene model prediction, we identified 5,434 nearly full-length genes, indicating the utility of draft genomes of *Protium* in our efforts to improve the understanding of TPS diversity in the genus. In the comparative analysis run with four rosids, the number of clusters unique of *P. kleinii* (1,795) was similar to those of the soya (2,120) and rice (2,035) genomes. This is not surprising since the angiosperm group is highly diversified [33] and it is the first time that the genome composition of a species of the Protium genus has been analyzed. On the other hand, these results suggest that the gene dataset of *P. kleinii* may be overestimated due to the splitting of the allelic variation in separated loci [31]. We cannot discard the possibility of recent gene duplication events [34]. Comparative analysis reported 26,497 singletons which can be the result of duplicated genes that evolved under positive selection [35]. However, they could also be artifacts generated during the step of gene prediction [36]. Last but not least, 1,300 clusters had at least one sequence from all of the other five species, but none for *P. kleinii*. This suggests that they could not be predicted with the applied methodology or they are probably absent in its genome. The availability of new genomes – especially genomes of “non-model” plants that produce a large diversity of terpenes – could facilitate the search for new compounds of interest or increase the production of known ones [9]. Furthermore, an improved knowledge base about plant terpene metabolism will facilitate the manipulation of the terpene pathway to change agronomic traits such as fruit flavors, floral scent, and plant defense against insects, and provide knowledge to engineer terpenoid production on a large-scale [37]. Although terpene compounds have diverse chemical structures, all are derived from two five-carbon isomeric basic building blocks: isopentenyldiphosphate (IPP) and dimethylallyldiphosphate (DMAPP) [38, 39]. In plants, production of IPP and DMAPP involves two pathways: the MVA and MEP pathways. The simultaneous condensation of IPP and DMAPP in various combinations gives rise to different intermediate precursors for the biosynthesis of plant terpenes. For more details see Tholl, 2015. Our chemical analysis of *P. kleinii* essential oil revealed the presence of the monoterpenes terpinolene (79.66%); limonene (8.72%); *α*-pinene (5.55%) and -phellandrene (1.57%). Monoterpenes are produced through many steps of oxidation and ionization of the GPP, which are catalyzed by monooxygenases and TPS, respectively, [37] and are the principal components of the volatile fraction of oleoresins from Protium species [25]. Terpinolene was the major component found in the essential oil from resin. This compound purportedly has therapeutic properties such as antifungal, antibacterial, anticancer, antioxidant and sedative activity [40, 41] and is also the most abundant compound of fresh *P. heptaphyllum* oleoresin [5]. Parts of the *P. heptaphyllum* genome were previously sequenced in order to study the repertoire of monoterpene synthases and to better understand monoterpene synthesis in Protium; they concluded that the chemical composition of *P. heptaphyllum* oleoresins changes over time [5]. We searched in our draft genome for genes involved in the pathways of terpene biosynthesis, which include the terpenoid backbone biosynthesis pathway and three other pathways which generate mono-, di-, tri- and sesquiterpenes. Our draft genome presents 68 genes – of which 20 are full length genes – involved in these terpene pathways. The presence of genes coding for enzymes involved in terpene biosynthesis shows that *P. kleinii* most likely produces the monoterpenes limonene, geraniol, *α*-terpineol, 1-8 cineole and (+)-neomenthol; the diterpene (E,E)-4,8,12-trimethyltrideca-1,3,7,11-tetraene (TMTT); the triterpene *β*-amyrin; and the sesquiterpenes (E,E)-farnesol, (E,E)-*α*-farnesene, (-)-germacrene D. In our essential oil chemical analysis we did not encounter certain monoterpenes that are reported in the literature as being produced by other *Protium* species, such as *α*-terpineol, 1,8-cineole and (+)-neomenthol [25]. However, the KEGG-mapped enzymes strongly suggest that *P. kleinii* has the terpene synthases that are responsible for the production of these compounds. Although geraniol is not mentioned in the literature about Protium resins, we mapped geranyl diphosphate diphosphatase, a key enzyme involved in its synthesis, and showed that geraniol probably can be produced by the plant, as observed with other rosids, such as Citrus [42] and Vitis [43]. Geraniol is an important alcohol present in essential oil from aromatic plants, and is largely used as a fragrance in industry. Geraniol is also reported to be present in flowers and vegetative tissues [44] and it is being studied as an alternative biofuel produced by microbial engineering [45]. The TPS and monooxygenases gene families – two main enzyme families involved in terpene synthesis – are molecularly highly diverse [37], which can jeopardize the identification of orthologues in the genomes of related species when conducting similarities searches. In fact, the terpene pathway is not fully identified in the genomes of four other rosids that we compared. It is important to note that it was possible to identify genes related to the production of the enzyme involved in *β*-amirin synthesis. *β*-amirin and *α*-amirin are reported as important components of the fixed fraction of the oleoresin *P. kleinii* [24, 46]. This is consistent with our chemical analysis of the volatile fraction, in which we, likewise, did not detect the presence of *β*-amirin or *α*-amirin. In this first draft genome, we could not find the synthases genes responsible for the TPS which, according to the KEGG database, produce the major compounds terpinolene, *α*-pinene and *α*-phellanderene. Exonerate did not succeed in finding them, perhaps because of the high molecular diversity within the *Protium* group [9] and in other plants [8] and the lack of a closer orthologue. Another possible explanation is the high fragmentation of the presented draft genome. Thus, the TPS genes responsible for the biosynthesis of the four major metabolites present in the volatile fraction of the *P. kleinii* resin were screened using the HMMsearch software. The phylogenetic tree shows that of the nine TPS queries of monoterpene pathways, only the limonene synthase of C. sinensis nested with sequences of *P. kleinii* in a well-supported clade. These data show that these enzymes may perform the same function in P. *kleinii*. With this analysis we also observed that the TPS repertoire in the genome of P. kleinii may be more extensive than what is described for other plants in this group [5, 9]. This may enable the discovery and study of new compounds of scientific and commercial interest. As terpenes extraction is laborious, can result in low yields, and is usually less cost-effective than chemical synthesis, biotechnological production represents an intriguing alternative, that may permit high regio- and stereoselectivity reactions and large-scale production [37, 47]

## 5. CONCLUSION

We generated a draft *P. Kleini*i genome and gene annotation that enabled us to predict 5,434 full length genes. Among these are 20 terpene synthesis pathways genes which code for specific terpene synthase (TPS) enzymes essential for the production of mono- and other terpenes. These data will directly support further studies to elucidate the molecular bases of these key aroma compounds, and will set the stage for future conservation research, bioprospecting, and novel biotechnology approaches to highly-scaled production of naturally synthesized terpene chemicals with important pharmaceutical and cosmetic applications.

## 6. METHODS

### A. Plant material

Samples of young leaves of *Protium kleinii* Cautrec. (Burseraceae), popularly known as *almécega* or white-tar, were collected at the Legado das Águas Reserve, in São Paulo, in southeastern Brazil, and deposited at the São Paulo municipal herbarium under the number PMSP 18043. Access to the Brazilian biodiversity was registered in the National System of Genetic Resource Management and Associated Traditional Knowledge under the number A0BCB8D.

### B. DNA Extraction, Genome Sequencing and Assembly

Genomic DNA was extracted from a young leaf of *P. kleinii* (10 cm2) using the DNeasy plant mini Kit^®^ (Qiagen, Hilden, Germany) following the manufacturer’s instructions. Sequencing was performed on an IlluminaHiSeq 2500 platform using 100 bp paired-end reads. The quality of the reads was analyzed by FastQC [48]. Low quality reads (<33 quality score) and below 80 bp in length were filtered using Trimmomatic [49]. K-mer analysis was then performed with Jellyfish [50] and genomescope browser [51]. Next, the reads were assembled *de novo* using SOAPdenovo2 [52]. After genome assembly, only scaffolds larger than 500 bp were maintained for further analysis. Repeats were *ab initio* identified with RepeatModeler [53] and the genome was masked using this *de novo* library of repeats and RepeatMasker[54]. Fastq files are available in Data S1.

### C. Assembly gene content assessment

In order to evaluate the completeness of the current genome assembly, we submitted all masked scaffolds to the gVolante server [55]. We set 1 as value for cut-off length, which means that all scaffolds were analyzed. Single-copy genes were predicted using AUGUSTUS [56] and then were assessed with lineage-specific hidden Markov profiles of Benchmarking Universal Single-Copy Orthologs (BUSCO) [57] in order to assign them to one of the orthologues groups.

### D. Gene Prediction and Annotation

Gene prediction was performed using the masked genome as input for AUGUSTUS (ver. 2.5.5; [56] trained for *A. thaliana* with parameters –genemodel=complete –strand=both – noInFrameStop=true. The functional annotation of the predicted proteins was determined through similarity and homology analysis using BLASTp (ver.2.5.0; [58] with a maximum E-value of 1e-05, in which the sequences were first searched against the UniProtKB/Swiss-Prot database [59]. All sequences that could not be identified at this stage were then submitted to BLASTp against the NCBI non-redundant database [60]. An automatic annotation of BLASTp results was carried out using Blast2GO (ver. 5.1.1; [61]). Metabolic pathway and orthology-oriented functional annotation of the protein sequences were performed against the KEGG database [21] using the KAAS [14] with the bi-directional best hit strategy and all datasets of dicot and monocot plants selected as reference organisms. All putative proteins were also annotated against all databases available in InterProScan (ver. 5.29-68.0; [62]) for conserved domains search and GO terms mapping.

### E. Comparative genomics

In order to find orthologues of *P. kleinii* shared with other species of plants, we performed a clustering analysis with OrthoVenn server [63] using all amino acid sequences deduced in the genome of *P. kleinii*, four rosids (*A. thaliana*, *C. sinensis*, *G. max* and *V. vinifera*; [15–18] and one monocotyledone (*O. sativa*; [19], as the outgroup. The datasets of proteins of each species available in OrthoVenn server were accessed in the Ensembl Plants repository [64], except for *C. sinensis* which was downloaded from the NCBI (ID: 10702; [15] and then manually uploaded. We set an E-value of 1e-5 as cutoff for all-to-all protein similarity comparisons and 1.5 as inflation value for the generation of orthologues clusters using the Markov Cluster Algorithm. The first protein added to the cluster was submitted to a BLASTp search against the UniProt database [59] and the first hit was used to infer the GO term assigned to one of the three categories (Biological process, molecular function, or cellular component). The plain text output containing all clusters was accessed and parsed with in-house scripts in order to identify orthologues shared with the other five species of plants, specific gene expansions of *P. kleinii*, and singletons.

### F. Essential oil analysis

Chemical analysis of the essential oil from 100 grams of *P. kleinii* resin was performed. The volatile fraction of the resin was separated by hydrodistillation in a Clevenger apparatus, dried with sodium sulfate anhydrous (Na2SO4), and then analyzed by GC-MS. Next, the oil was diluted 1:100 in hexane (Sigma Aldrich, Brazil) and the analyses were made on a Shimadzu QP2010 Plus system using a DB5-MS fused silica capillary column (30 m, 0.25 mm I.D., 0.25 μm film thickness). The oven temperature was programmed to increase 3° C per minute over the course of one hour from 60° C to 240° C where the temperature was maintained for another 50 minutes. Other operating conditions were: carrier gas: He (99.999%); flow rate: 1.03 mL/min; injector temperature: 250° C; interface temperature: 250° C and ion source temperature: 200° C. Samples were injected in the split mode (1:50). Mass spectra were obtained under electroionization at 70 eV. Mass range: m/z 40 D to 500 D. The identification of individual components was based on comparison of mass spectral fragmentation patterns with those available in the NIST Mass Spectral Library and the GC Kovats retention Indices (KI) on a DB-5 column and literature data [65].

### G. Terpene Biosynthesis Pathways genes

All genes belonging to terpene biosynthesis pathways were recovered from the KAAS result using the KO numbers as reference. This was done using in-house scripts. In order to confirm the presence and absence of the orthologues of *P. kleinii* in the monoterpene biosynthesis pathway, we searched for them in all soft masked scaffolds using Exonerate software [66]. Next, we used the amino acid sequence of the closest orthologue of *P. kleinii* available in KEGG database as a query [Additional Table 11]. All coding sequences found were translated using Transeq software [67] and submitted to BlastKOALA [21] in order to confirm whether one or more of the retrieved sequences belonged to the same KO group of the respective query. Additionally, the orthology relationship between of *P. kleinii* genes and other TPS [Additional Table 11 and fragment 2 described in Zapata and Fine [9]] was inferred through phylogenetic analysis. For this purpose, we first searched all members of the family within the *P. kleinii* protein dataset with HMMsearch [68] using the respective conserved domain profile available in the Protein Family Database (Pfam; [22]. All TPS were aligned with MAFFT [69], then the global alignment was manually inspected with Seaview [70] and used to build a neighbor-joining tree using MEGA DNA version 7 [71] with 500 replicates of bootstrap [72]. When proteins grouped together in clades with 80% or higher bootstrap support, we marked them as candidate genes that were highly likely to belong to the family of terpene biosynthesis pathways.

## 7. AVAILABILITY OF DATA AND MATERIALS

Compressed files comprising the Fastq obtained with the illumina run can be accessed in **our database**, however, the access might be granted through the corresponding author permission.

## 10. ACKNOWLEDGEMENTS

Authors are indebted to Americo JA for critical review, Mrs Lisboa M and dr Fonseca MF for their support. Parabotanist Martins Flores MJ and Dr. Zandona L for invaluable help in field activities. Jankovic N for performing chemical analysis of the essential oil.

## REFERENCES

1. Lange BM. The Evolution of Plant Secretory Structures and Emergence of Terpenoid Chemical Diversity. Annu Rev Plant Biol. 2015;66:139–59. doi:10.1146/annurev-arplant-043014-114639.

2. Siedenburg G, Jendrossek D, Breuer M, Juhl B, Pleiss J, Seitz M, et al. Activation-independent cyclization of monoterpenoids. Appl Environ Microbiol. 2012;78:1055–62. doi:10.1128/AEM.07059-11.

3. Bution ML, Molina G, Abrahaõ MRE, Pastore GM. Genetic and metabolic engineering of microorganisms for the development of new flavor compounds from terpenic substrates. Crit Rev Biotechnol. 2015;35:313–25. doi:10.3109/07388551.2013.855161.

4. Paramasivan K, Mutturi S. Progress in terpene synthesis strategies through engineering of Saccharomyces cerevisiae. Crit Rev Biotechnol. 2017;37:974–89. doi:10.1080/07388551.2017.1299679.

5. Albino RC, Oliveira PC, Prosdocimi F, da Silva OF, Bizzo HR, Gama PE, et al. Oxidation of monoterpenes in Protium heptaphyllum oleoresins. Phytochemistry. 2017;136:141–6. doi:10.1016/j.phytochem.2017.01.013.

6. da Silva ER, Oliveira DR de, Fernandes PD, Bizzo HR, Leitão SG. Ethnopharmacological Evaluation of Breu Essential Oils from Protium Species Administered by Inhalation. Evid Based Complement Alternat Med. 2017;2017 March 2012:1–10. doi:10.1155/2017/2924171.

7. Bohlmann J, Meyer-Gauen G, Croteau R. Plant terpenoid synthases: Molecular biology and phylogenetic analysis. Proceedings of the National Academy of Sciences. 1998;95:4126–33. doi:10.1073/pnas.95.8.4126.

8. Chen F, Tholl D, Bohlmann J, Pichersky E. The family of terpene synthases in plants: A mid-size family of genes for specialized metabolism that is highly diversified throughout the kingdom. Plant J. 2011;66:212–29. doi:10.1111/j.1365-313X.2011.04520.x.

9. Zapata F, Fine PVA. Diversification of the monoterpene synthase gene family (TPSb) in Protium, a highly diverse genus of tropical trees. Mol Phylogenet Evol. 2013;68:432–42. doi:10.1016/j.ympev.2013.04.024.

10. Davis EM, Croteau R. Cyclization Enzymes in the Biosynthesis of Monoterpenes, Sesquiterpenes, and Diterpenes. In: F J L, J C V, editors. Topics in Current Chemistry. Springer, Berlin, Heidelberg; 2000. p. 53–95.

11. Uliano-Silva M, Dondero F, Dan Otto T, Costa I, Lima NCB, Americo JA, et al. A hybrid-hierarchical genome assembly strategy to sequence the invasive golden mussel, Limnoperna fortunei. Gigascience. 2018;7. doi:10.1093/gigascience/gix128.

12. Singh S, Singh DB, Singh S, Shukla R, Ramteke PW, Misra K. Exploring Medicinal Plant Legacy for Drug Discovery in Post-genomic Era. Proceedings of the National Academy of Sciences, India Section B: Biological Sciences. 2018. doi:10.1007/s40011-018-1013-x.

13. trinityrnaseq. Github. https://github.com/trinityrnaseq/trinityrnaseq. Accessed 20 Sep 2018.

14. Moriya Y, Itoh M, Okuda S, Yoshizawa AC, Kanehisa M. KAAS: An automatic genome annotation and pathway reconstruction server. Nucleic Acids Res. 2007;35:182–5. doi:10.1093/nar/gkm321.

15. Xu Q, Chen LL, Ruan X, Chen D, Zhu A, Chen C, et al. The draft genome of sweet orange (Citrus sinensis). Nat Genet. 2013;45:59–66. doi:10.1038/ng.2472.

16. Swarbreck D, Wilks C, Lamesch P, Berardini TZ, Garcia-Hernandez M, Foerster H, et al. The Arabidopsis Information Resource (TAIR): gene structure and function annotation. Nucleic Acids Res. 2008;36 Database issue:D1009–14. doi:10.1093/nar/gkm965.

17. Schmutz J, Cannon SB, Schlueter J, Ma J, Mitros T, Nelson W, et al. Genome sequence of the palaeopolyploid soybean. Nature. 2010;463:178. doi:10.1038/nature08670.

18. Jaillon O, Aury J-M, Noel B, Policriti A, Clepet C, Casagrande A, et al. The grapevine genome sequence suggests ancestral hexaploidization in major angiosperm phyla. Nature. 2007;449:463. doi:10.1038/nature06148.

19. Mahesh HB, Shirke MD, Singh S, Rajamani A, Hittalmani S, Wang G-L, et al. Indica rice genome assembly, annotation and mining of blast disease resistance genes. BMC Genomics. 2016;17:242. doi:10.1186/s12864-016-2523-7.

20. Bausher MG, Singh ND, Lee SB, Jansen RK, Daniell H. The complete chloroplast genome sequence of *Citrus sinensis* (L.) Osbeck var “Ridge Pineapple”: Organization and phylogenetic relationships to other angiosperms. BMC Plant Biol. 2006;6:1–11. doi:10.1186/1471-2229-6-21.

21. Kanehisa M, Sato Y, Morishima K. BlastKOALA and GhostKOALA: KEGG Tools for Functional Characterization of Genome and Metagenome Sequences. J Mol Biol. 2016;428:726–31. doi:10.1016/j.jmb.2015.11.006.

22. Finn RD, Coggill P, Eberhardt RY, Eddy SR, Mistry J, Mitchell AL, et al. The Pfam protein families database: towards a more sustainable future. Nucleic Acids Res. 2016;44:279–85. doi:10.1093/nar/gkv1344.

23. Mukherjee S, Stamatis D, Bertsch J, Ovchinnikova G, Verezemska O, Isbandi M, et al. Genomes OnLine Database (GOLD) v.6: data updates and feature enhancements. Nucleic Acids Res. 2017;45:D446–56. doi:10.1093/nar/gkw992.

24. Otuki MF, Vieira-Lima F, Malheiros, Yunes RA, Calixto JB. Topical antiinflammatory effects of the ether extract from *Protium kleinii* and *α*-amyrin pentacyclic triterpene. Eur J Pharmacol. 2005;507:253–9. doi:10.1016/j.ejphar.2004.11.012.

25. Rüdiger AL, Siani AC, Veiga Junior VF. The chemistry and pharmacology of the South America genus *Protium* Burm. f. (Burseraceae). Pharmacogn Rev. 2007;1:93–104.

26. Zappi DC, Ranzato Filardi FL, Leitman P, Souza VC, Walter BMT, Pirani JR, et al. Growing knowledge: An overview of Seed Plant diversity in Brazil. Rodriguesia. 2015;66:1085–113. doi:10.1590/2175-7860201566411.

27. Wang X, Xu Y, Zhang S, Cao L, Huang Y, Cheng J, et al. Genomic analyses of primitive, wild and cultivated citrus provide insights into asexual reproduction. Nat Genet. 2017;49:765–72. doi:10.1038/ng.3839.

28. Yandell M, Ence D. A beginner’s guide to eukaryotic genome annotation. Nat Rev Genet. 2012;13:329–42. doi:10.1038/nrg3174.

29. Shimizu T, Tanizawa Y, Mochizuki T, Nagasaki H, Yoshioka T, Toyoda A, et al. Draft sequencing of the heterozygous diploid genome of satsuma (*Citrus unshiu* Marc.) using a hybrid assembly approach. Front Genet. 2017;8 DEC:1–19. doi:10.3389/fgene.2017.00180.

30. The Arabidopsis Genome Initiative. Analysis of the genome sequence of the flowering plant *Arabidopsis thaliana*. Nature. 2000;408:796. doi:10.1038/35048692.

31. Denton JF, Lugo-Martinez J, Tucker AE, Schrider DR, Warren WC, Hahn MW. Extensive error in the number of genes inferred from draft genome assemblies. PLoS Comput Biol. 2014;10:e1003998. doi:10.1371/journal.pcbi.1003998.

32. Veeckman E, Ruttink T, Vandepoele K. Are We There Yet? Reliably Estimating the Completeness of Plant Genome Sequences. 2016;28 August:1759–68. doi:10.1105/tpc.16.00349.

33. Judd WS, Campbell CS, Kellogg EA, Stevens PF. Plant systematics a phylogenetic approach. Sinauer; 2015.

34. Kaessmann H, Vinckenbosch N, Long M. RNA-based gene duplication: mechanistic and evolutionary insights. Nat Rev Genet. 2009;10:19. doi:10.1038/nrg2487.

35. Kaessmann H. Origins, evolution and phenotypic impact of new genes. Genome Res. 2010;:gr – 101386. doi:10.1101/gr.101386.109.

36. Prabh N, Rödelsperger C. Are orphan genes protein-coding, prediction artifacts, or non-coding RNAs? BMC Bioinformatics. 2016;17:226. doi:10.1186/s12859-016-1102-x.

37. Pateraki I, Heskes AM, Hamberger B. Cytochromes P450 for terpene functionalisation and metabolic engineering. In: Schrader J, Bohlmann J, editors. Biotechnology of Isoprenoids. Springer; 2015. p. 107–39. doi:10.1007/102014301.

38. Trapp SC, Croteau RB. Genomic organization of plant terpene synthases and molecular evolutionary implications. Genetics. 2001;158:811–32.

39. Tholl D. Biosynthesis and biological functions of terpenoids in plants. In: Biotechnology of isoprenoids. Springer; 2015. p. 63–106.

40. Okumura N, Yoshida H, Nishimura Y, Kitagishi Y, Matsuda S. Terpinolene, a component of herbal sage, downregulates AKT1 expression in K562 cells. Oncol Lett. 2012;3:321–4. doi:10.3892/ol.2011.491.

41. Aydin E, Türkez H, Taşdemir S. Anticancer and antioxidant properties of terpinolene in rat brain cells. Archives of Industrial Hygiene and Toxicology. 2013;64:415–24. doi:10.2478/10004-1254-64-2013-2365.

42. Gancel A-L, Ollitrault P, Froelicher Y, Tomi F, Jacquemond C, Luro F, et al. Leaf volatile compounds of six citrus somatic allotetraploid hybrids originating from various combinations of lime, lemon, citron, sweet orange, and grapefruit. J Agric Food Chem. 2005;53:2224–30. doi:10.1021/jf048315b.

43. Martin DM, Aubourg S, Schouwey MB, Daviet L, Schalk M, Toub O, et al. Functional annotation, genome organization and phylogeny of the grapevine (Vitis vinifera) terpene synthase gene family based on genome assembly, FLcDNA cloning, and enzyme assays. BMC Plant Biol. 2010;10:226.

44. Chen W, Viljoen AM. Geraniol—a review of a commercially important fragrance material. S Afr J Bot. 2010;76:643–51. doi:10.1016/j.sajb.2010.05.008.

45. Liu W, Xu X, Zhang R, Cheng T, Cao Y, Li X, et al. Engineering Escherichia coli for high-yield geraniol production with biotransformation of geranyl acetate to geraniol under fed-batch culture. Biotechnol Biofuels. 2016;9:58. doi:10.1186/s13068-016-0466-5.

46. Lima FV, Malheiros A, Otuki MF, Calixto JB, Yunes RA, Cechinel Filho V, et al. Three new triterpenes from the resinous bark of *Protium kleinii* and their antinociceptive activity. J Braz Chem Soc. 2005;163 B:578–82. doi:10.1590/s0103-50532005000400014.

47. Schempp FM, Drummond L, Buchhaupt M, Schrader J. Microbial Cell Factories for the Production of Terpenoid Flavor and Fragrance Compounds. J Agric Food Chem. 2018;66:2247–58. doi:10.1021/acs.jafc.7b00473.

48. Babraham Bioinformatics - FastQC A Quality Control tool for High Throughput Sequence Data. http://www.bioinformatics.babraham.ac.uk/projects/fastqc/. Accessed 20 Sep 2018.

49. Bolger AM, Lohse M, Usadel B. Trimmomatic: A flexible trimmer for Illumina sequence data. Bioinformatics. 2014;30:2114–20. doi:10.1093/bioinformatics/btu170.

50. Margais G, Kingsford C. A fast, lock-free approach for efficient parallel counting of occurrences of k-mers. Bioinformatics. 2011;27:764–70. doi:10.1093/bioinformatics/btr011.

51. Vurture GW, Sedlazeck FJ, Nattestad M, Underwood CJ, Fang H, Gurtowski J, et al. GenomeScope: fast reference-free genome profiling from short reads. Bioinformatics. 2017;33:2202–4. doi:10.1093/bioinformatics/btx153.

52. Luo R, Liu B, Xie Y, Li Z, Huang W, Yuan J, et al. F1000Prime recommendation of: SOAPdenovo2: an empirically improved memory-efficient short-read de novo assembler. Gigascience. 2012; July:1–6. doi:10.1186/2047-217X-1-18.

53. Smit AFA, Hubley R. RepeatModeler Open-1.0. Available fom http://wwwrepeatmaskerorg.2016.

54. Smit AFA, Hubley R, Green P. RepeatMasker Open-4.0. Available fom http://wwwrepeatmaskerorg.2016. http://www.repeatmasker.org.

55. Nishimura O, Hara Y, Kuraku S. GVolante for standardizing completeness assessment of genome and transcriptome assemblies. Bioinformatics. 2017;33:3635–7. doi:10.1093/bioinformatics/btx445.

56. Hoff KJ, Stanke M. WebAUGUSTUS–a web service for training AUGUSTUS and predicting genes in eukaryotes. Nucleic Acids Res. 2013;41 Web Server issue:123–8. doi:10.1093/nar/gkt418.

57. Simão FA, Waterhouse RM, Ioannidis P, Kriventseva EV, Zdobnov EM. BUSCO: Assessing genome assembly and annotation completeness with single-copy orthologs. Bioinformatics. 2015;31:3210–2. doi:10.1093/bioinformatics/btv351.

58. Altschul SF, Madden TL, Schäffer AA, Zhang J, Zhang Z, Miller W, et al. Gapped BLAST and PSI-BLAST: a new generation of protein database search programs. Nucleic Acids Res. 1997;25:3389–402. doi:10.1093/nar/25.17.3389.

59. UniProt Consortium. UniProt: a hub for protein information. Nucleic Acids Res. 2014;43:D204–12.

60. Benson DA, Cavanaugh M, Clark K, Karsch-Mizrachi I, Lipman DJ, Ostell J, et al. GenBank. Nucleic Acids Res. 2017;45:D37–42. doi:10.1093/nar/gkw1070.

61. Conesa A, Götz S, García-Gómez JM, Terol J, Talón M, Robles M. Blast2GO: a universal tool for annotation, visualization and analysis in functional genomics research. Bioinformatics. 2005;21:3674–6. doi:10.1093/bioinformatics/bti610.

62. Jones P, Binns D, Chang HY, Fraser M, Li W, McAnulla C, et al. InterProScan 5: Genome-scale protein function classification. Bioinformatics. 2014;30:1236–40. doi:10.1093/bioinformatics/btu031.

63. Wang Y, Coleman-Derr D, Chen G, Gu YQ. OrthoVenn: a web server for genome wide comparison and annotation of orthologous clusters across multiple species. Nucleic Acids Res. 2015;43:W78–84. doi:10.1093/nar/gkv487.

64. Yates A, Akanni W, Amode MR, Barrell D, Billis K, Carvalho-Silva D, et al. Ensembl 2016. Nucleic Acids Res. 2016;44:D710–6. doi:10.1093/nar/gkv1157.

65. Adams RP. Identification of Essential Oil Components by Gas Chromatography. 41st edition. Allured publishing; 2017.

66. Slater GSC, Birney E. Automated generation of heuristics for biological sequence comparison. BMC Bioinformatics. 2005;6:31. doi:10.1186/1471-2105-6-31.

67. Rice P, Longden I, Bleasby A. EMBOSS: the European molecular biology open software suite. Trends Genet. 2000;16:276–7. doi:10.1016/S0168-9525(00)02024-2.

68. Eddy SR. Accelerated profile HMM searches. PLoS Comput Biol. 2011;7:e1002195. doi:10.1371/journal.pcbi.1002195.

69. Katoh K, Rozewicki J, Yamada KD. MAFFT online service: multiple sequence alignment, interactive sequence choice and visualization. Brief Bioinform. 2017. doi:10.1093/bib/bbx108.

70. Gouy M, Guindon S, Gascuel O. SeaView version 4: a multiplatform graphical user interface for sequence alignment and phylogenetic tree building. Mol Biol Evol. 2010;27:221–4. doi:10.1093/molbev/msp259.

71. Kumar S, Stecher G, Tamura K. MEGA7: molecular evolutionary genetics analysis version 7.0 for bigger datasets. Mol Biol Evol. 2016;33:1870–4. doi:10.1093/molbev/msw054.

72. Efron B, Halloran E, Holmes S. Bootstrap confidence levels for phylogenetic trees. Proceedings of the National Academy of Sciences. 1996;93:13429.

